# Adaptation to Environmental Variability Shapes Dormancy in *Daphnia*

**DOI:** 10.64898/2026.05.06.723256

**Authors:** Robert J. Porter, Lauren A. Bradshaw, Isabelle F. Marsh, Madison E. Doceti, Alan O. Bergland

## Abstract

Dormancy is a widespread adaptive strategy that allows organisms to survive in temporally varying habitats by suspending development and reproduction. Although environmental variability is expected to shape dormancy strategies, it remains unclear how differences in environmental variability and predictability influence both the production of dormant embryos and the termination of dormancy. We addressed these questions by comparing *D. pulex* and *D. obtusa*, two closely related species that inhabit environments differing in variability and predictability. We hypothesized that *D. obtusa*, which inhabits ephemeral environments, would exhibit a greater propensity for sexual reproduction and dormancy and would require stronger cues to break dormancy than D. *pulex*, which occurs in more permanent, predictable habitats. Consistent with our hypothesis, *D. obtusa* lineages produced significantly more males and ephippia than *D. pulex* when reared under identical laboratory conditions, indicating greater investment in sexual reproduction and dormancy. Contrary to our hypothesis, we found no difference in responsiveness to cues between the two species. Across species, embryos broke dormancy and hatched most readily after experiencing changes in cold and light, even if not experienced at the same time. In contrast, desiccation reduced the propensity to break dormancy. Together, these results indicate that species occupying more ephemeral environments invest more heavily in the production of dormant offspring, but that the environmental cues regulating dormancy termination appear broadly similar between species. This pattern suggests that while investment in dormancy may evolve in response to environmental variability, the mechanisms controlling dormancy termination are more conserved.

## Introduction

Dormancy is a common strategy that enables organisms to withstand temporally variable environments (Cohen, 1966; Shoemaker & Lennon, 2018). A key factor shaping the evolution of dormancy strategies is the predictability of environmental change. Predictability can reflect both how consistently unfavorable conditions occur, and how precisely their timing can be inferred from environmental cues (ten Brink et al., 2020). In some systems, environmental deterioration is predictable in both occurrence and timing. For example, when habitat suitability is driven by regular seasonal changes, organisms can use reliable token cues such as photoperiod, to closely align development with favorable conditions (Grevstad & Coop, 2015). In contrast, many organisms inhabit more variable, ephemeral environments, where habitat deterioration is inevitable, yet the precise timing of when conditions become inhospitable remains uncertain. As a result, environmental cues provide only coarse information about future conditions, making it difficult for organisms to precisely time the initiation or termination of dormancy. Uncertainty in the end of the growing period is expected to favor alternative strategies, such as continuous or diversified investment in dormant offspring, which reduces the risk of complete reproductive failure when conditions deteriorate unpredictably (Pires et al., 2023; Wourms, 1972). Although the effects of environmental predictability and variability on dormancy strategies have been widely studied, relatively few studies directly compare dormancy strategies across systems that experience strongly contrasting levels of environmental variability, and even fewer examine both the initiation and termination of dormancy.

Freshwater habitats provide an excellent opportunity to study how species adapt to variation in habitat quality. Permanent lakes and ephemeral pools differ fundamentally in their hydroperiods, the duration and timing of water availability, which shapes the reliability of environmental cues as signals of favorable conditions (Brendonck et al., 2017; De Meester et al., 2005; Rosbakh et al., 2020). In permanent freshwater environments, cues such as photoperiod and seasonal temperature changes can be reliable indicators of habitat change (Bernhardt et al., 2020). In ephemeral environments, where environmental change occurs much more rapidly, organisms more heavily rely on dormancy to cope with unpredictable rainfall and water loss (Schröder et al., 2007). The interplay between environmental variability and dormancy strategy makes freshwater habitats useful for examining how species adapt to temporal environmental change.

Among freshwater taxa, *Daphnia* are a particularly well-suited system to study the evolution of dormancy strategies. *Daphnia spp.* can be found in a diverse range of environments, from permanent lakes to ephemeral rock pools (Altermatt & Ebert, 2010). During favorable conditions, *Daphnia* predominantly reproduce clonally (Barnard-Kubow et al., 2022; Mikulski & Grzesiuk, 2020). When conditions begin to deteriorate, *Daphnia* increase the production of males, who then mate with females to produce dormant sexual resting embryos in a case called an ephippium (Gerber et al, 2018). Ephippia serve as the dispersal stage of *Daphnia* and can survive harsh environmental conditions, remaining viable across seasons (Pietrzak & Ślusarczyk, 2006). *Daphnia* ephippia usually contain up to two sexually produced embryos depending on mating success (Winsor & Innes, 2002). An ephippium is produced in place of an asexual clutch, which would typically contain 2-40 offspring (Gliwicz & Lampert, 1994), representing a shift from rapid clonal population growth to the production of fewer, more durable sexual offspring. Dormant offspring then generally require a change in environmental cues to terminate dormancy and emerge (Radzikowski et al., 2018). However, a subset of ephippial offspring can continue development and hatch without any change in environmental cues, emerging into the same environment in which they were produced (Cáceres, 1998; Porter et al., 2023). Although environmental cues regulating dormancy in *Daphnia* have been well studied, fewer studies have examined how variation in ephippia production and dormancy termination together shape life history strategies.

This variation in dormancy strategies is especially informative when examined across closely related taxa. The *D. pulex* clade provides such a system, as the clade is made up of several sister species, including *D. pulex*, *D. obtusa*, and *D. pulicaria* (Ye et al., 2022). *D. pulex* and *D. obtusa* are thought to have diverged ∼31 million years ago (Murray et al., 2025) and, while these species are morphologically similar, they can generally be differentiated based on specific diagnostic traits (Schwartz et al., 1985). *D. obtusa* typically inhabit more ephemeral environments, such as ephemeral pools (Nix & Jenkins, 2000), whereas *D. pulex* can be found in a broader range of habitats, from ephemeral pools to permanent ponds and lakes (Barnard-Kubow et al., 2022; Wen Deng, 1997). Within species, individuals can also differ in their propensity to enter dormancy. *Daphnia* from more permanent environments tend to invest less in dormancy and invest more heavily in clonal reproduction (Tessier & Cáceres, 2004) and can hatch even in the absence of changing external cues (Porter et al., 2023). In contrast, *Daphnia* from ephemeral habitats tend to produce more males and ephippia, reflecting a greater reliance on dormancy as a survival strategy (Altermatt & Ebert, 2010; Barnard-Kubow et al., 2022).

To investigate the evolution of dormancy strategies, we examined how dormancy initiation and termination vary within and between these two species that inhabit environments that differ in the extent and predictability of growing periods. Specifically, we asked whether patterns of dormant offspring production vary between the species, whether offspring from each species can exit dormancy in the absence of external cues, and how environmental conditions shape the rate of dormancy termination. First, we hypothesized that the *D. obtusa,* from the more ephemeral environment, will have a higher propensity to invest in dormancy, and produce more males and ephippia than *D. pulex* when grown in similar conditions. Second, we hypothesized that dormant offspring collected from *D. pulex*, sourced from larger, deeper ponds, which are more predictable and less variable, will terminate dormancy early in the absence of external cues. Additionally, we hypothesized that desiccation would trigger dormancy termination more readily in *D. obtusa*, whereas *D. pulex* would be more responsive to changing temperatures. Overall, our results show that *Daphnia* lineages from ephemeral habitats invest more heavily in dormancy, producing more males and resting eggs than lineages from more permanent environments. However, we observed no difference between species in their responsiveness to environmental cues that terminate dormancy.

## Methods

### Study Populations

We collected *D. obtusa* and *D. pulex* across England during March of 2023 (Fig. 1A, Table S1). Each geographic site had multiple ponds located at a relatively close geographic distance. Habitat quality was assessed using multiple indicators, including estimates of pond ephemerality (estimated from depth measurements, <= 0.6m) and pH (Fig. 1B). Live *Daphnia* samples were transported back to the University of Virginia, where we established clonal lineages from single female isolates. We maintained clonal lineages at 18°C and 16H:8H light:dark (L:D) conditions in 250 ml glass jars, where the *Daphnia* underwent asexual reproduction. *Daphnia* lineages were maintained in hard artificial pond water (ASTM) (Standard, 2007) and were fed *Chlorella vulgaris* purchased from Reed Mariculture (*Chlorella vulgaris* Superfresh V12), which was centrifuged and resuspended in ASTM for feeding.

**Fig. 1:**
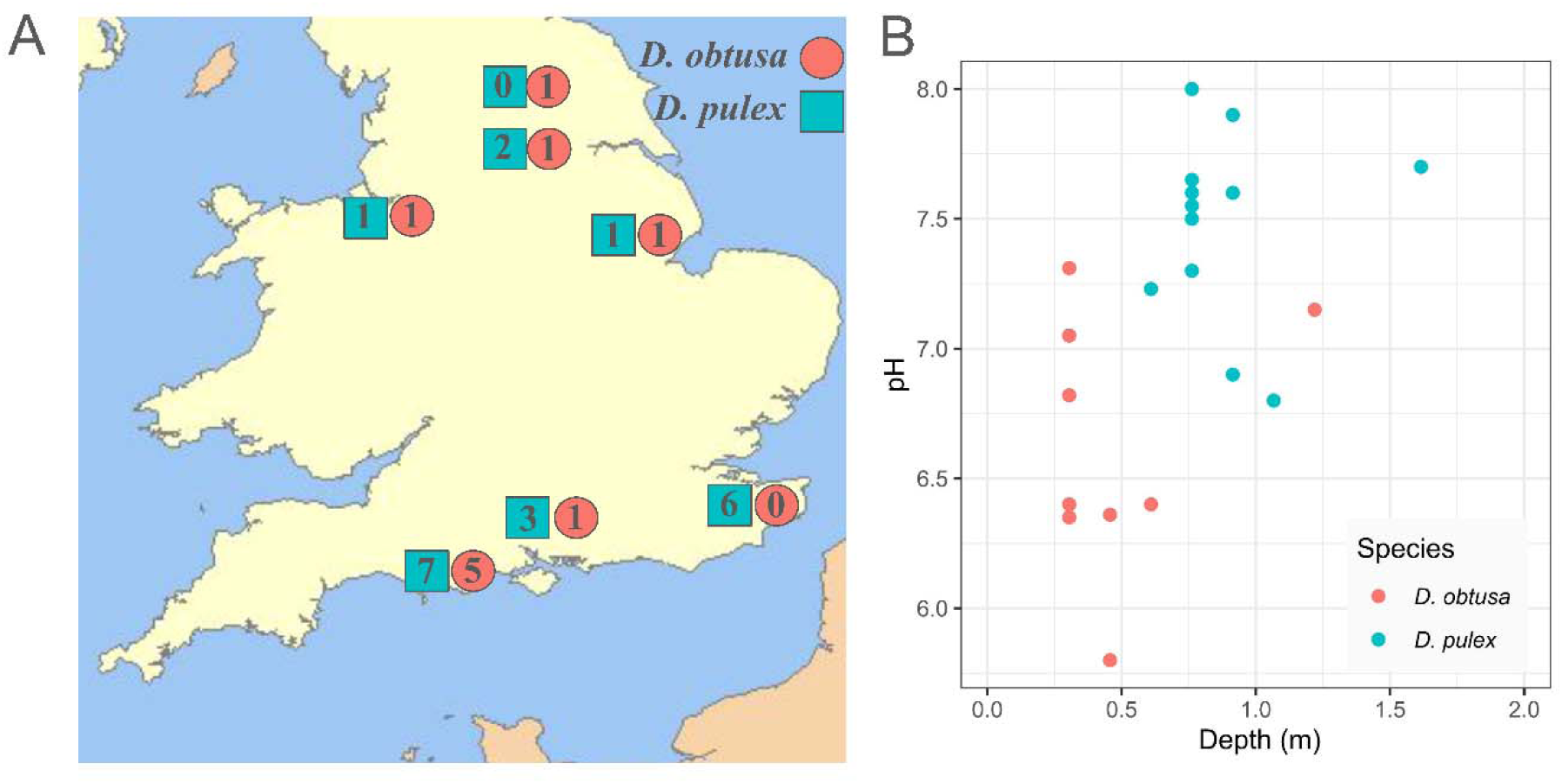
Sampling locations and habitat characteristics of *D. obtusa* and *D. pulex* ponds across England. (A) Map of pond locations. Numbers indicate how many ponds are in each location, and colors indicate the dominant species in each pond. (B): Depth vs Water pH of each pond, colored by dominant species of each pond. *D. obtusa* are found in shallower, more acidic environments. The *D. obtusa* pond that had a depth of ∼1.2m was a roadside ditch sampled immediately after rain had occurred.

### Species identification and mitochondrial phylogenetic analysis

We morphologically identified individuals to the *pulex* clade, which includes *D. pulex* and *D. obtusa (Schwartz et al., 1985).* Additionally, we included a single *D. ambigua* individual from North America (Belle Isle, Pool 759, 37.5265N, -77.4565W) to serve as an outgroup for subsequent analyses. From the lab lines, we isolated a subset of individuals to undergo treatment for DNA extraction. We placed ∼30 individuals from each lineage into a Sephadex (Sephadex Superfine-G25) and antibiotic treatment (streptomycin, tetracycline, and ampicillin, 50 mg/l of each) for 2 full days to clear their digestive tract and reduce associated microbiota. Individuals were then preserved in 70% ethanol for sequencing. For lineages that did not survive the antibiotic treatment, we used ∼10 individuals from previously untreated ethanol samples. As the *D. ambigua* individual was field collected and preserved in ethanol, it did not undergo antibiotic treatment. After preservation, ∼10 individuals were homogenized using a TissueLyser II (30 hz for 20 seconds, flipped then repeated), and RNA was removed using RNase-A. We extracted genomic DNA using an Axygen AxyPrep MAG Tissue Genomic DNA prep kit, and we constructed libraries using a scaled down Nextera protocol (Baym et al., 2015). We size selected the libraries using a BluePippin to 300-500bp, and quantified size and concentration using a TapeStation high sensitivity D1000 kit. Libraries were sequenced using PE150bp reads on a Novaseq X. We then trimmed adapter sequences from the reads using Trimmomatic ver. 0.39 (Bolger et al., 2014).

To species-type individuals, we also generated a cytochrome oxidase I (COI) reference panel using COI gene sequences from 23 *Daphnia* species/lineages (STable 2). Trimmed reads were mapped to the reference panel using bwa/0.1.17 (Li, 2013). We used Samtools/1.17 (Li et al., 2009) to count the number of reads that mapped to COI genes from each species. For each library, we then normalized the species-specific counts by calculating the proportion of reads mapping to each species’ COI gene relative to the total reads mapped across all COI references. Lineages with fewer than 40 mapped reads were removed, and only those with at least 80% of normalized reads mapping to a single species were retained. These assignments were used to determine the species identity of each individual in subsequent analyses.

Trimmed sequences were then mapped to their respective mitochondrial reference genome. For each species, we generated VCF files from the resulting BAM files, and filtered SNPs for a depth greater than 20, and QUAL greater than 20. We generated consensus sequences from those VCF’s back onto their reference genome using bcftools/v1.17 consensus (Danecek et al., 2021). We then used BEAUti to convert the fasta file into a .xml file, and phylogenetic analyses were conducted in BEAST/v2.7.7 (Bouckaert et al., 2019). A maximum clade credibility tree was generated in Treeannotator, which we then pruned down to only our UK samples, as well as a single North American *D. ambigua* to root. In R/v4.4.1 (R Core, 2021), we used dplyr/v1.1.4 (Wickham et al, 2023) and ape/v5.8.1 (Paradis & Schliep 2019) to combine clone metadata to the tree, which we then plotted using ggtree/v3.12.0 (Yu et al., 2017; Yu et al., 2018; Yu, 2020; Xu et al., 2022; Yu, 2022).

### Mesocosm rearing

We reared 28 lineages (two biological replicates each, totaling 56 mesocosms) for eight weeks under a semi-randomized block design, with replicates run concurrently at 18C, under a long day (16L:8D) photoperiod. We seeded the mesocosms with ∼20 adult females in 2L of media, doubling the media every two days until mesocosms reached 16L. We fed the mesocosms with 95,000 cells/mL *C. vulgaris* every Monday, Wednesday, and Friday for the first five weeks, and increased the concentration of *C. vulgaris* to 142,000 cells/mL after week 5 due to increased population density. We sampled mesocosms weekly for 8 weeks to collect ephippia, and preserved a subset of individuals at week 5 in ethanol to determine age and demography. To sample the mesocosms, we first stirred them to mix the contents, then removed 1 L of media and collected all individuals in that sample before replacing the volume with fresh media.

### Ephippia collection and hatching assay

We collected ephippia on a weekly basis from each mesocosm and stored ephippia individually in clear 96 well plates (N=12,417 ephippia). Weekly sampling was conducted to minimize the likelihood of ephippia hatching prior to collection. We filled wells with ASTM and placed the plates in 18°C under long day conditions (16L:8D) for four weeks in the same Environmental Growth Chamber they were produced in. We checked plates weekly using a Leica S8 APO microscope to determine hatch rate. The microscopes lights were adjusted to be as dim as possible, but still maintain visualization. After a month of observation for early hatching, plates were placed into one of four conditions: Cold/Wet/Dark, Cold/Dry/Dark, Warm/Wet/Light, and Warm/Dry/Light. Cold temperatures were set at 4°C, while warm temperatures were 18°C. For the Dry treatment, ephippia were desiccated by maintaining plates uncovered in a growth chamber, while ephippia in the Wet treatment were left hydrated. Dark was continuous darkness, while the Light treatment was long day conditions (16L:8D). After removing the plates from the conditions back into 18°C long day, we measured hatching for two more weeks to determine the hatch rate.

To assess investment in dormant offspring, we estimated the total number of embryos produced per lineage. After all observable hatching ceased, which took about two weeks, we treated a subset of ephippia (N=5,144 ephippia) with a 2% sodium hypochlorite solution for 10-15 minutes (Paes et al., 2016) to visualize the embryos and determine the fill rate, defined as the total number of embryos divided by the maximum possible number of embryos (assuming 2 embryos per ephippia). We then estimated the total number of individuals by summing the early-hatched, late-hatched, and unhatched embryos per lineage within each replicate.

### Experimental test of dormancy cues

To further investigate the environmental factors that trigger the emergence from dormancy, we conducted an experiment using a single lineage of *D. obtusa* (P19_50). We used this lineage because it produced many ephippia, allowing us to test different combinations of environmental conditions on the emergence from dormancy. We collected 1,533 ephippia from a similar mesocosm experiment as described above, and monitored them for a month to measure early hatching. To more specifically determine the cues related to hatching, we conducted a fully factorial experiment under three environmental conditions. These included changing temperature (4C vs 18C), photoperiod (16L:8D vs complete darkness), and desiccation status (dried vs wet), with ∼192 ephippia per treatment. Ephippia were placed under these conditions for four weeks, after which individuals were rehydrated and moved back into warm, long-day wet conditions to assess hatching. We then tracked ephippia for two more weeks to determine how many individuals hatched.

### Data analysis

We analyzed reproductive traits and population size using generalized linear mixed models in R (V4.41, R Core Team, 2021), choosing the error structure based on the type of response variable. Traits expressed as proportions, including the proportion of males in the population, ephippial fill rate, and the proportion of embryos hatching early or late, were analyzed using models with a binomial error distribution (Models 2, 3, 5, 6, 7 - see below). When traits were measured as counts, such as the number of individuals, or the number of ephippia produced, we used a Posson error distribution (Models 1 & 4 - see below).

Across analyses, we accounted for the hierarchical structure of the data by including clonal identity and experimental replicate as random effects. Replicates were nested within clonal identity to reflect multiple measurements from the same clonal lineage. The general form of the mixed effect models was:

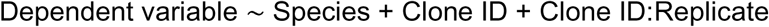

To test for differences between species, we compared models with and without species as a fixed effect using a Likelihood Ratio Test (Chambers & Hastie, 1992). To determine whether clonal lineages within a species differed from one another, we repeated these analyses separately for each species. Within each species, we compared models that included Clone ID (clonal lineage) as a random effect to models that only accounted for variation among replicates. A significant improvement in model fit indicated that clonal lineages varied, explaining additional variation beyond differences among replicates.

To examine factors that influenced late hatching, we modeled the proportion of embryos that hatched late as a function of species, temperature (Cold/Warm), and desiccation treatment (Dry/Wet), including all interaction terms. Replicates nested within clonal id were included to account for repeated measurements from the same lineage. The model structure was:

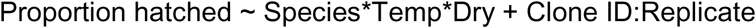

To analyze hatching in the fully factorial experiment using only *D. obtusa*, we modeled the proportion of embryos that hatched as a function of temperature (Cold/Warm), desiccation status (Dry/Wet), and light regime (Dark/Long Day), along with their interactions. The model structure was:

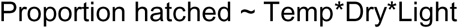

Statistical analyses were conducted in R (v4.4.1; R Core Team, 2021) using the following packages: data.table/v1.16.0 (Barrette et al., 2025), ggplot2/v3.5.2 (Wickham, 2016), foreach/v1.5.2 (Microsoft & Weston, 2022), lattice/v0.22-6 (Sarkar, 2008), tidyr/v1.3.1 (Wickham et al., 2024), tidyverse/v2.0.0 (Wickham et al., 2019), gridExtra/v2.3 (Auguie, 2017), dplyr/v1.1.4 (Wickham, 2023), reshape2/v1.4.4 (Wickham, 2007), patchwork/v1.3.1 (Pedersen, 2025), knitr/v1.50 (Xie, 2014, 2015, & 2025), lme4/v1.1-37 (Bates et al, 2015), cowplot/v1.2.0 (Wilke, 2025), and ggsignif/v0.6.4 (Constantin & Patil, 2021).

## Results

We tested whether species occupying habitats with different levels of environmental variability differ in reproductive investment and dormancy-related traits. Using mesocosm experiments with multiple clonal lineages of *Daphnia obtusa* and *Daphnia pulex*, we quantified population density, male production, ephippia production, and ephippial fill rates. We then compared these traits between species and among clonal lineages.

### Species typing using the COI gene

We sequenced 93 lineages originating from 35 ponds. Of the 82 lineages that yielded usable COI sequence data, 29 were identified as *D. obtusa* and 53 as *D. pulex* based on BLAST comparisons to reference sequences (Fig. 2). Eleven sequenced lineages were excluded from further analyses due to insufficient read depth or failure to recover high-quality COI sequence. We additionally included 23 published COI reference sequences to anchor species assignments; however, none of our samples used in this study showed ambiguous placement relative to these references. Most lineages used in subsequent experiments were successfully sequenced. For the small number that did not have sequence coverage to determine species identity (n=11), species identity was inferred from morphology and from the species assignments of other sequenced individuals from the same pond.

**Fig. 2:**
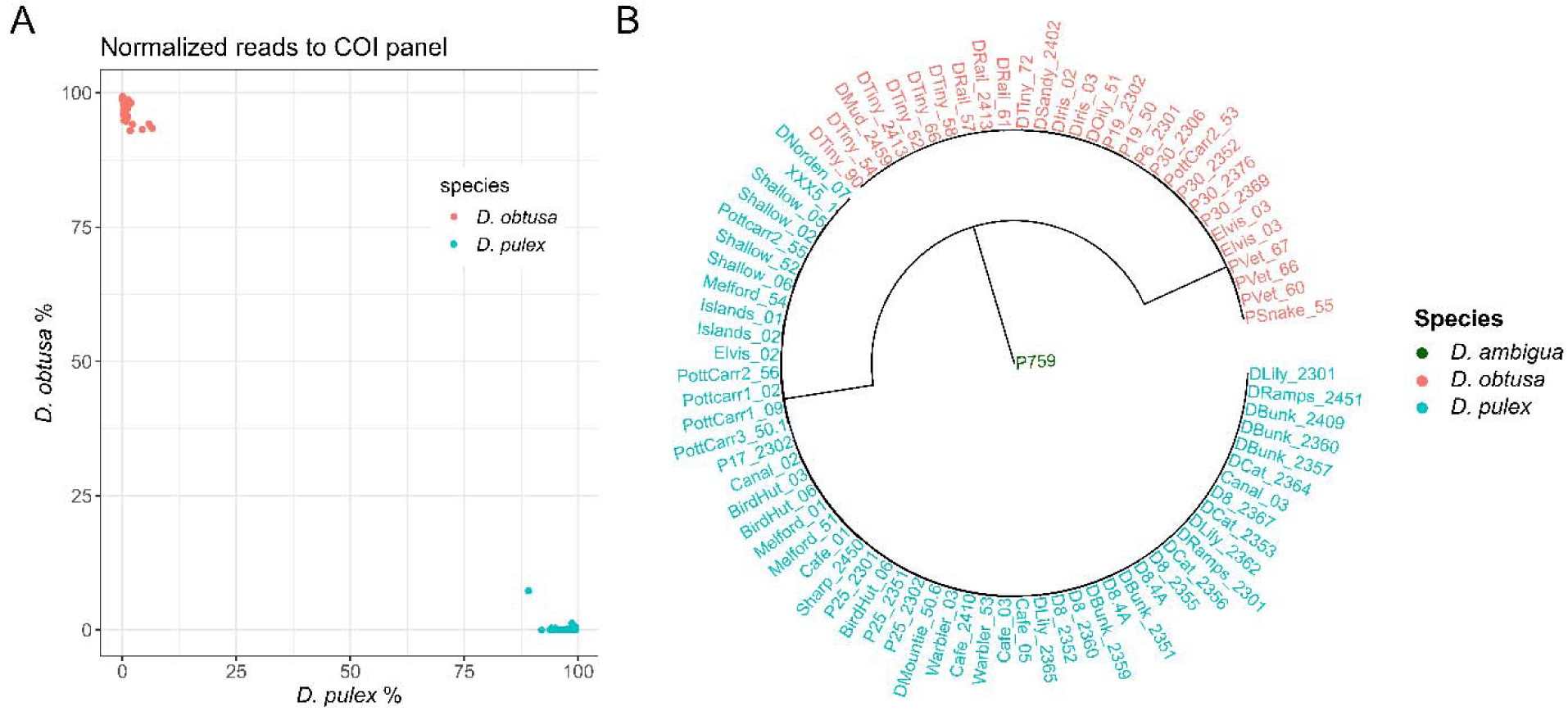
Species identification and phylogenetic placement of samples based on mitochondrial COI sequences. (A) Mapping of raw reads to a COI reference panel. Samples were filtered to >40 reads in at least one reference species. *D. obtusa* and *D. pulex* each group separately. (B) Phylogeny of samples that passed species-typing, rooted to North American *D. ambigua*. *D. obtusa* and *D. pulex* group separately. Only the ponds Pottcarr2 and Elvis had both species observed from sequencing.

### Ephippia and male production

In the mesocosms, we found no difference in population density between *D. obtusa* or *D. pulex* (Fig. 3A, *χ*²=0.19, p=0.89, df=1, Table 1 - Model 1A). Clone ID did not significantly affect the model fit for either *D. obtusa* (Fig. 3A’, *χ*²=0.047, df=1, p=0.83, Table 1 - Model 1B) or *D. pulex* (Fig. 3A’, *χ*²=1.41, df=1, p=0.24, Model 1C), while accounting for replicate. This suggests that there was not consistent variation in population density attributable to genetic differences among clones.

**Fig. 3:**
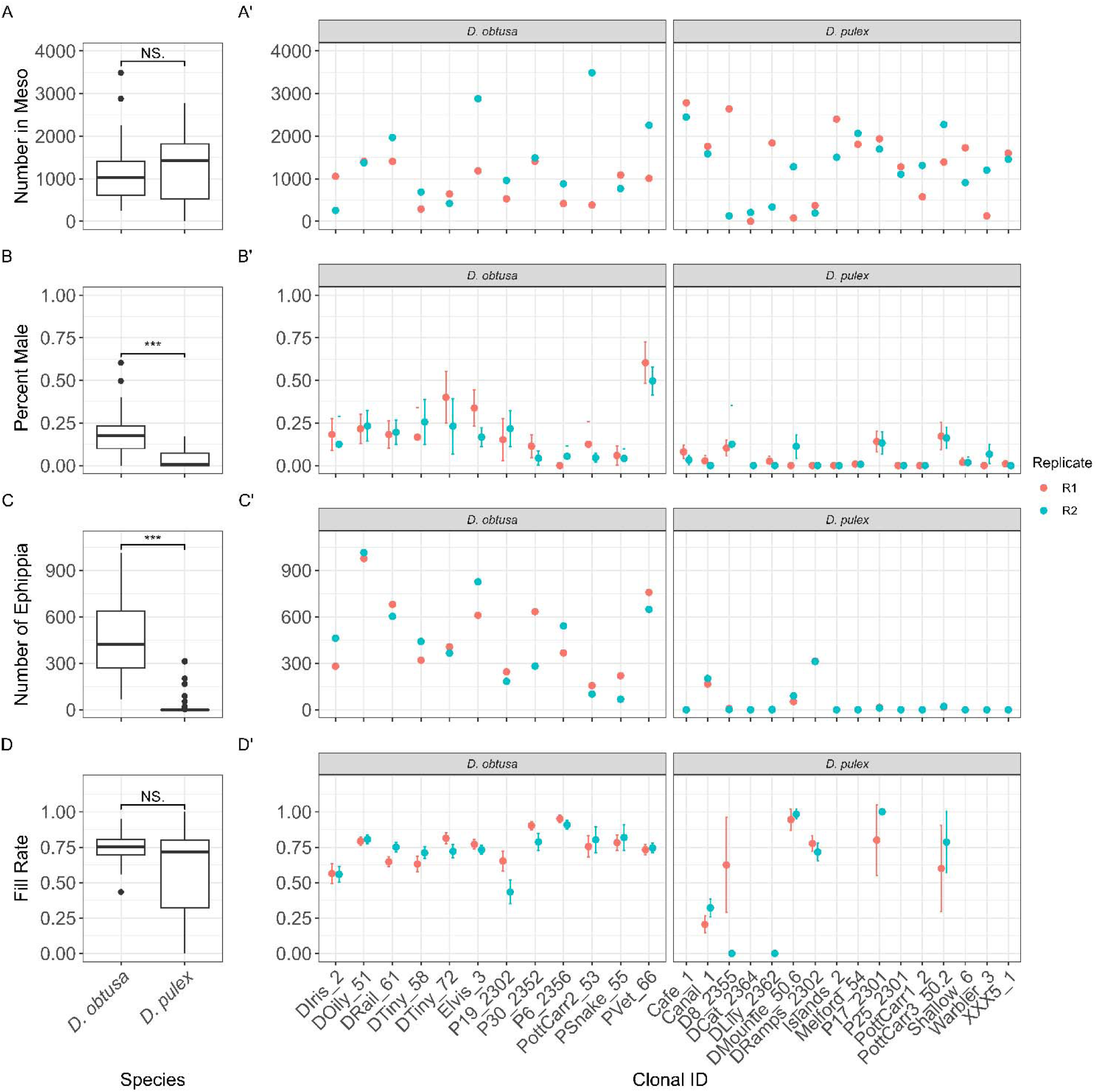
Population density and reproductive traits of *D. obtusa* and *D. pulex* in mesocosms. (A) Population density did not differ between species, and there was not clonal variation within species (A′). (B) *D. obtusa* produced a higher percent of males than *D. pulex*, with significant clonal variation in both species (B′). (C) *D. obtusa* lineages produced more ephippia, and Clone ID significantly improved model fit for both species (C′). (D) Average ephippial fill rate did not differ between, but varied significantly among clones within each species (D′). Significance stars indicate p < 0.05 (*), p < 0.01 (**), and p < 0.001 (***) from linear mixed-model.

**Table 1:** Models 1 to 5.

|  | Comparison | Fixed Effects | Random Effects | npar | AIC | BIC | logLik | Deviance | Chisq | df | P-value | Significance |
| --- | --- | --- | --- | --- | --- | --- | --- | --- | --- | --- | --- | --- |
| Model 1A | Population estimates at week 5 (*D. obtusa* vs *D. pulex*) | Species | Clone ID + Clone ID:Replicate | 4 | 615.5 | 623.6 | -303.75 | 0.0187 | 0.0187 | 1 | 0.8913 |  |
| Model 1B | Population estimates at week 5 (*D. obtusa*) | None | Clone ID + Clone ID:Replicate | 3 | 251.43 | 254.96 | -122.71 | 0.0474 | 0.0474 | 1 | 0.8276 |  |
| Model 1C | Population estimates at week 5 (*D. pulex*) | None | Clone ID + Clone ID:Replicate | 3 | 362.34 | 366.74 | -178.17 | 1.4105 | 1.4105 | 1 | 0.235 |  |
| Model 2A | Total ephippia produced (*D. obtusa* vs *D. pulex*) | Species | Clone ID + Clone ID:Replicate | 4 | 525.55 | 534.82 | -258.77 | 23.309 | 23.309 | 1 | 1.38E-06 | *** |
| Model 2B | Ephippia production (*D. obtusa*) | None | Clone ID + Clone ID:Replicate | 3 | 331.68 | 335.22 | -162.84 | 9.6225 | 9.6225 | 1 | 0.001922 | ** |
| Model 2C | Ephippia production (*D. pulex*) | None | Clone ID + Clone ID:Replicate | 3 | 164.03 | 169.83 | -79.016 | 33.668 | 33.668 | 1 | 6.54E-09 | *** |
| Model 3A | Male production at week 5 (*D. obtusa* vs *D. pulex*) | Species | Clone ID + Clone ID:Replicate | 4 | 282.93 | 290.95 | -137.46 | 14.528 | 14.528 | 1 | 0.000138 | *** |
| Model 3B | Male production (*D. obtusa*) | None | Clone ID + Clone ID:Replicate | 3 | 157.87 | 161.41 | -75.937 | 10.76 | 10.76 | 1 | 0.001037 | ** |
| Model 3C | Male production (*D. pulex*) | None | Clone ID + Clone ID:Replicate | 3 | 125.74 | 130.04 | -59.87 | 14.509 | 14.509 | 1 | 0.00014 | *** |
| Model 4A | Fill rate of ephippia (*D. obtusa* vs *D. pulex*) | Species | Clone ID + Clone ID:Replicate | 4 | 319.95 | 326.4 | -155.98 | 0.4904 | 0.4904 | 1 | 0.4838 |  |
| Model 4B | Fill rate (*D. obtusa*) | None | Clone ID + Clone ID:Replicate | 3 | 236.51 | 240.04 | -115.25 | 9.4111 | 9.4111 | 1 | 0.002157 | ** |
| Model 4C | Fill rate (*D. pulex*) | None | Clone ID + Clone ID:Replicate | 3 | 81.833 | 83.528 | -37.917 | 7.2134 | 7.2134 | 1 | 0.007236 | ** |
| Model 5A | Early hatching (*D. obtusa* vs *D. pulex*) | Species | Clone ID + Clone ID:Replicate | 4 | 125.83 | 132.05 | -58.915 | 0.8312 | 0.8312 | 1 | 0.3619 |  |
| Model 5B | Early hatching (*D. obtusa* lineages) | None | Clone ID + Clone ID:Replicate | 3 | 97.561 | 101.09 | -45.78 | 6.4065 | 6.4065 | 1 | 0.01137 | * |
| Model 5C | Early hatching (*D. pulex* lineages) | None | Clone ID + Clone ID:Replicate | 3 | 28.889 | 30.083 | -11.445 | 1.9688 | 1.9688 | 1 | 0.1606 |  |

*D. obtusa* lineages produced a significantly higher proportion of males when compared to *D. pulex* (Fig. 3B, *χ*²=14.5, p=1.3x10⁻_, df=1, Table 1 - Model 2A). Including Clone ID as a factor significantly improved model fit for both *D. obtusa* (Fig. 3B’, *χ*²=10.8, df=1, p=0.0010, Table 1 - Model 2B) and *D. pulex* (Fig. 3B’, *χ*²=14.5, df=1, p=0.00014, Table 1 - Model 2C), demonstrating that there is genetic variation in male production rates between clonal lineages for both species.

Additionally, *D. obtusa* lineages produced significantly more ephippia (Fig. 3C, *χ*²=23.3, p=1.4x10⁻_, df=1, Table 1 - Model 3A) than *D. pulex*. Both the *D. obtusa* (Fig. 3C’, *χ*²=9.6, df=1, p=0.0019, Table 1 - Model 3B) and *D. pulex* (Fig. 3C’, *χ*²=33.7, df=1, p=6.5e-09, Table 1 - Model 3C) models significantly improved when adding Clone ID as a factor, similarly indicating that there is consistent differences in ephippia production among clonal lineages.

We found no differences in average ephippial fill rates between the two species (Fig. 3D, *χ*²=0.49, p=0.48, Table 1 - Model 4A). However, including Clone ID as a factor significantly improved model fit for both *D. obtusa* (Fig. 3D’, *χ*²=9.4, df=1, p=0.0022, Table 1 - Model 4B) and *D. pulex* (Fig. 3D’, *χ*²=7.2, df=1, p=0.0072, Table 1 - Model 4C). Because *D. obtusa* produced more ephippia overall, the similar fill rates between species indicate that *D. obtusa* lineages produced a greater total number of dormant embryos.

### Early Hatching

We observed a low rate of early hatching in both species. Early hatching in *D. obtusa* was around 0.31% (51 early hatchers out of 16,745 total embryos, 95% CI: 0.23%-0.40%). In *D. pulex*, early hatching was around 0.77% (10/1,296, 95% CI: 0.42%-1.41%). There was no significant difference in the proportion of early hatchers between species (Fig. 4A, *χ*²=0.83, df=1, p=0.36, Table 1 - Model 5A). There was significant variation among *D. obtusa* clonal lineages (Fig. 4B, *χ*²=6.4, df=1, p=0.011, Table 1 - Model 5B), but not among *D. pulex* lineages (Fig. 4B, *χ*²=2.0, df=1, p=0.16, Table 1 - Model 5C).

**Fig. 4:**
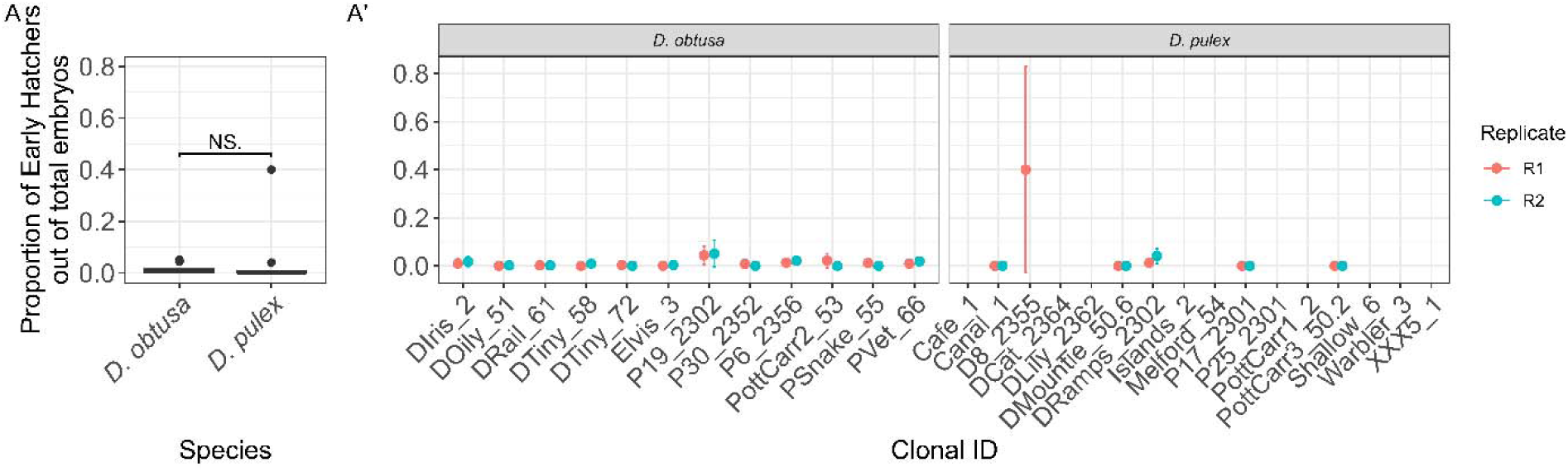
Rates of Early Hatching between *D. obtusa* and *D. pulex*. (A) Both species showed low rates of early hatching, with *D. obtusa* at 0.31% and *D. pulex* at 0.77%. There was no significant difference between species. (A′) Significant, but minor, between-clone variation was detected in *D. obtusa*, but not in *D. pulex*.

### Late hatching

Hatching from dormancy increased following exposure to environmental cues. An average of 13.3% of *D. obtusa* embryos (N = 2,232/16,745; 95% CI: 12.8-13.9%) and 11.7% of *D. pulex* embryos (N=152/1,296; 95% CI: 10.1-13.6%) emerged following a change in environmental cues. Among the four environmental treatments, the highest rates of late emergence occurred under cold (4°C), wet conditions, where 31.6% of *D. obtusa* embryos (N = 2,161/6,849; 95% CI: 30.5-32.7%) and 26.9% of *D. pulex* embryos (N = 145/539; 95% CI: 23.3-30.8%) hatched (Fig. 5A). The remaining three environmental treatments produced consistently low hatching rates. A generalized linear model revealed significant effects of species (*χ*² = 12.9, df = 1, p = 0.00033), temperature (*χ*² = 3984, df = 1, p < 2.2 × 10⁻¹_), desiccation (*χ*² = 35.4, df = 1, p = 2.67 × 10⁻_), and clonal identity (*χ*² = 536.8, df = 16, p < 2.2 × 10⁻¹_) on late hatching (Table 2 - Model 6). A strong interaction between temperature and desiccation was also detected (*χ*² = 61.3, df = 1, p = 4.78 × 10⁻¹_), indicating that the effect of temperature depended on moisture. There was also a significant degree of variation among clonal lineages in the Cold/Wet treatment for *D. obtusa* (*χ*² = 14.1, df = 1, p = 0.00018; Table 3 - Model 6B), but no significant variation was observed among *D. pulex* lineages under the same conditions (*χ*² = 1.79, df = 1, p = 0.18; Table 3 - Model 6C).

**Fig. 5:**
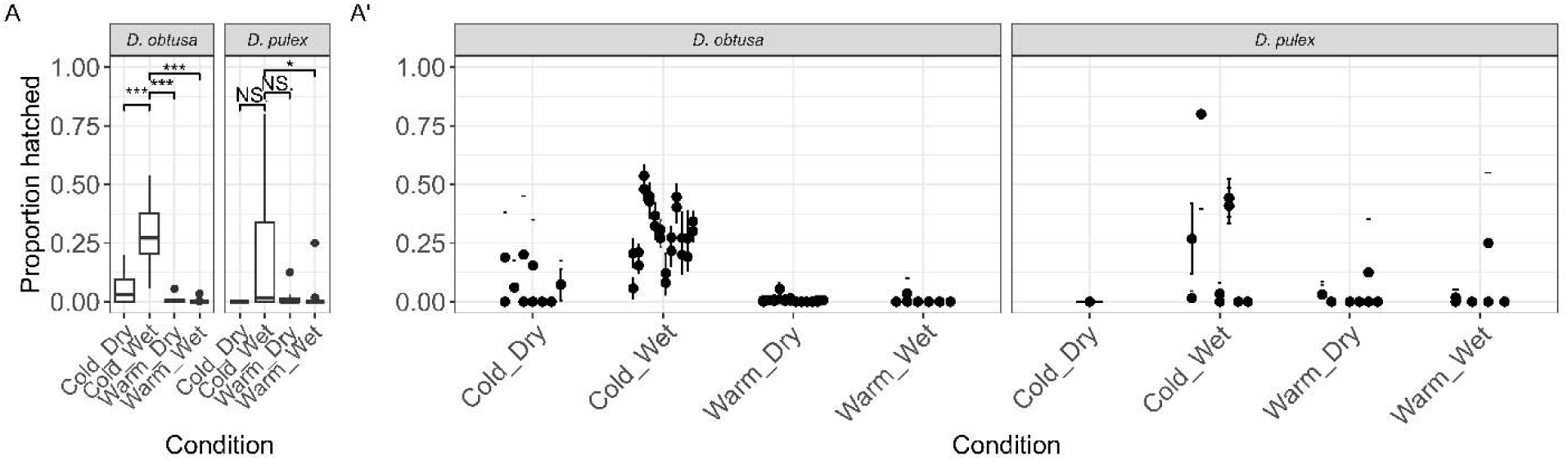
Late hatching responses of *D. obtusa* and *D. pulex* embryos under different environmental conditions. (A) Both species showed moderate levels of late hatching, with slightly higher rates in *D. obtusa* than in *D. pulex*. The highest hatching rates occurred under cold, wet conditions, where a substantial proportion of embryos from both species emerged. (A′) Significant between-clone variation was observed in *D. obtusa* under the Cold/Wet treatment, but not in *D. pulex*. The remaining environmental treatments had consistently low hatching rates.

**Table 2:** Model 6A.

|  |  |  |  |  |  |
| --- | --- | --- | --- | --- | --- |
| Model 6A | Late hatching (Temperature × Desiccation;<br>clone = "Clone ID") |  |  |  |  |
| Species model (with nested<br>random effects) |  |  |  |  |  |
| Effect | Df | Deviance<br>($\chi^2$) | Residual<br>Df | Residual<br>Dev. | p |
| - (Null) |  |  | 115 | 4829.5 |  |
| Species | 1 | 12.9 | 114 | 4816.6 | 0.00033 |
| Condition (Temp) | 1 | 3984 | 113 | 832.6 | <2.2e-16 |
| Dry (Desiccation) | 1 | 35.4 | 112 | 797.2 | 2.67e-09 |
| Clone ID | 16 | 536.8 | 96 | 260.4 | <2.2e-16 |
| Species × Condition | 1 | 0.1 | 95 | 260.4 | 0.799 |
| Species × Dry | 1 | 2.2 | 94 | 258.1 | 0.134 |
| Condition × Dry | 1 | 61.3 | 93 | 196.8 | 4.78e-15 |
| Clone ID : Replicate | 17 | 53.1 | 76 | 143.7 | 1.39e-05 |
| Species × Condition × Dry | 1 | 0 | 75 | 143.7 | 0.931 |

**Table 3:**
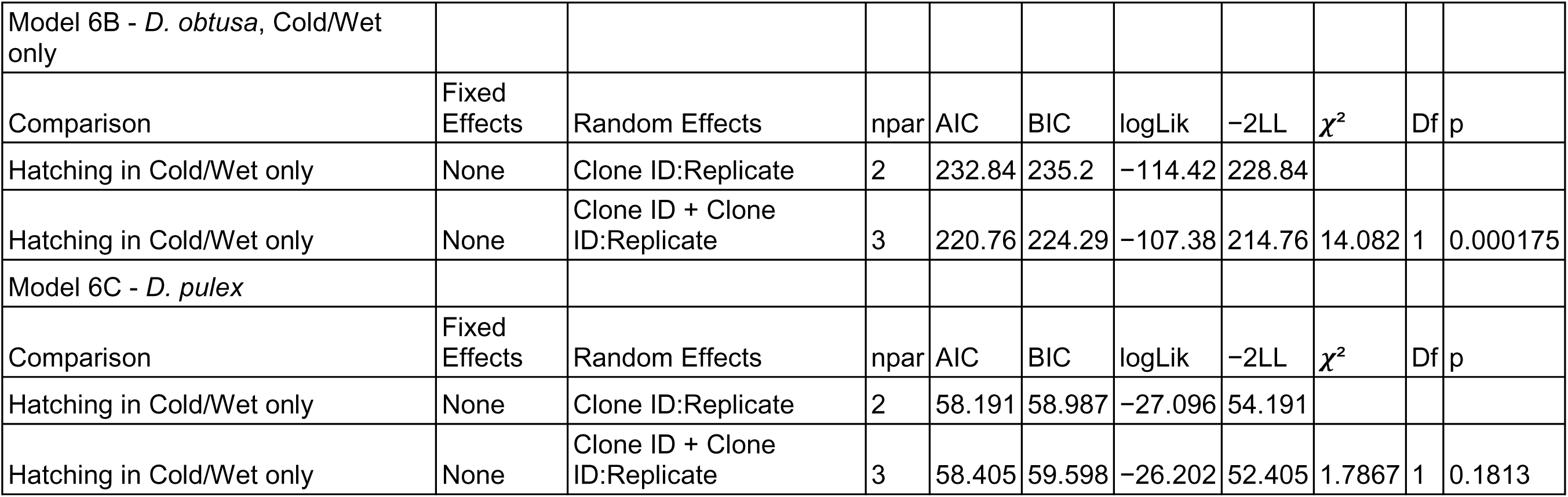
Models 6B and 6C.

|  |  |  |  |  |  |  |  |  |  |  |
| --- | --- | --- | --- | --- | --- | --- | --- | --- | --- | --- |
| Model 6B - *D. obtusa*, Cold/Wet only |  |  |  |  |  |  |  |  |  |  |
| Comparison | Fixed Effects | Random Effects | npar | AIC | BIC | logLik | -2LL | $\chi^2$ | Df | p |
| Hatching in Cold/Wet only | None | Clone ID:Replicate | 2 | 232.84 | 235.2 | -114.42 | 228.84 |  |  |  |
| Hatching in Cold/Wet only | None | Clone ID + Clone ID:Replicate | 3 | 220.76 | 224.29 | -107.38 | 214.76 | 14.082 | 1 | 0.000175 |
| Model 6C - *D. pulex* |  |  |  |  |  |  |  |  |  |  |
| Comparison | Fixed Effects | Random Effects | npar | AIC | BIC | logLik | -2LL | $\chi^2$ | Df | p |
| Hatching in Cold/Wet only | None | Clone ID:Replicate | 2 | 58.191 | 58.987 | -27.096 | 54.191 |  |  |  |
| Hatching in Cold/Wet only | None | Clone ID + Clone ID:Replicate | 3 | 58.405 | 59.598 | -26.202 | 52.405 | 1.7867 | 1 | 0.1813 |

### Hatching in response to different cues

Of the 1,533 ephippia (∼2,020 embryos) placed into the fully factorial hatching treatments, the highest hatching success occurred under Cold/Wet/Light conditions, where about three-quarters (75.5%, 95% CI [69.8, 80.4%]) of embryos hatched (190/252; Fig. 6). The next highest hatching was observed under Cold/Wet/Dark conditions, with around 43% [37.2, 49.3%] hatching (108 out of 250). In all other treatments, hatching remained low, averaging about 5.1% [4.1, 6.4%] overall (78/1,518; Fig. 6). There were strong effects of temperature (*χ*² = 265.58, df = 1, p < 2.2 × 10⁻¹_), desiccation (*χ*² = 256.70, df = 1, p < 2.2 × 10⁻¹_), and light (*χ*² = 18.10, df = 1, p = 2.10 × 10⁻_) on hatching success (Table 4 - Model 7). Significant interactions were also detected between temperature and desiccation (*χ*² = 136.88, df = 1, p < 2.2 × 10⁻¹_) and between temperature and light (*χ*² = 36.18, df = 1, p = 1.80 × 10⁻_), indicating that the effect of light and moisture depended on temperature conditions. The interaction between desiccation and light was not significant (*χ*² = 1.25, df = 1, p = 0.264). A significant three-way interaction among temperature, desiccation, and light was also detected (*χ*² = 10.03, df = 1, p = 0.00154), consistent with the particularly high hatching success observed under Cold/Wet/Light conditions.

**Fig. 6:**
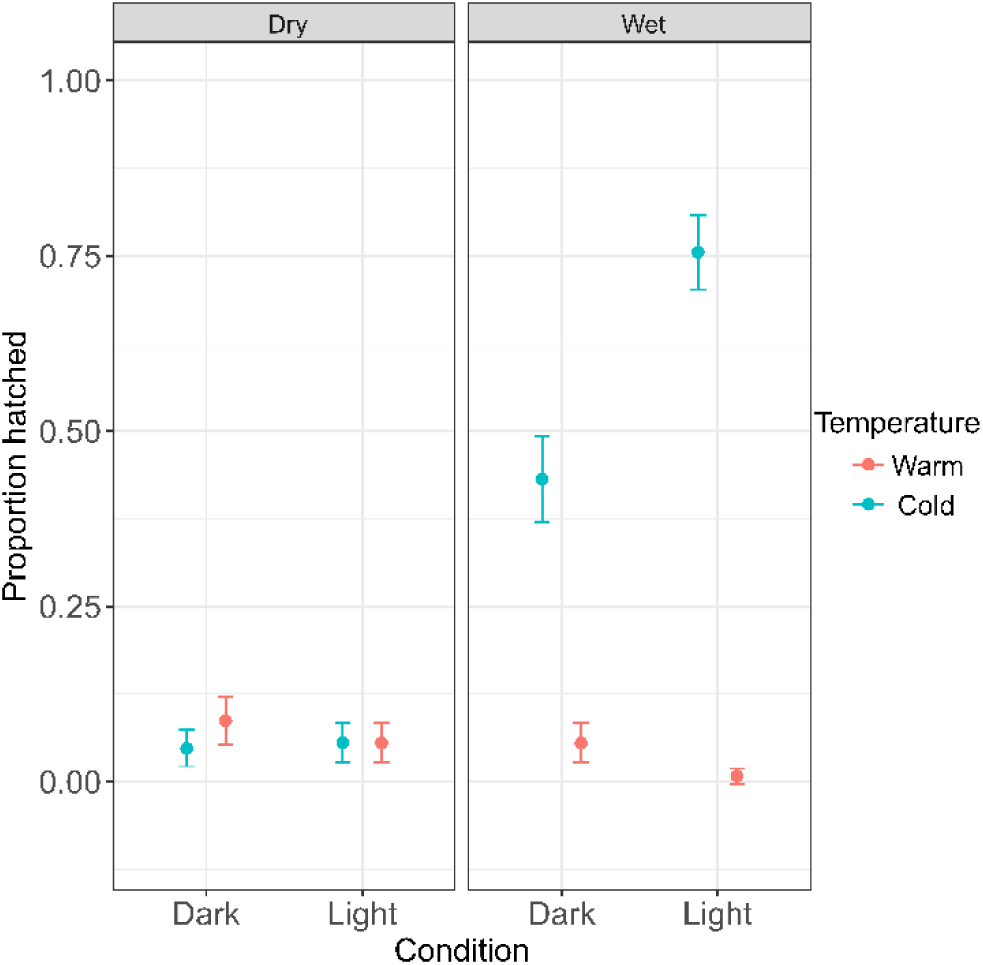
Factorial hatching in a lineage of *D. obtusa* comparing desiccation, light condition, and temperature. The highest hatching occurred in Cold/Wet/Light, followed by Cold/Wet/Dark. All other combinations of conditions had relatively low rates of hatching.

**Table 4:** Fully factorial hatching, Model 7.

|  |  |  |  |  |  |
| --- | --- | --- | --- | --- | --- |
| Model 7 - |  |  |  |  |  |
| Effect | Df | Deviance<br>($\chi^2$) | Residual Df | Residual<br>Dev. | p |
| (Null) |  |  | 7 | 724.71 |  |
| Temp | 1 | 265.578 | 6 | 459.13 | <2.2e-16 |
| Dried | 1 | 256.704 | 5 | 202.43 | <2.2e-16 |
| Light | 1 | 18.095 | 4 | 184.33 | 2.10e-05 |
| Temp × Dried | 1 | 136.884 | 3 | 47.45 | <2.2e-16 |
| Temp × Light | 1 | 36.178 | 2 | 11.27 | 1.80e-09 |
| Dried × Light | 1 | 1.246 | 1 | 10.03 | 0.264 |
| Temp × Dried × Light | 1 | 10.026 | 0 | 0 | 0.00154 |

## Discussion

In this study, we compared two closely related *Daphnia* species that occupy habitats that differ in environmental variability to examine how investment and emergence from dormancy varies across species and lineages. We hypothesized that *D. obtusa*, which is associated with more variable ephemeral habitats, would invest more heavily in sexual reproduction and resting egg production and exhibit broader responses to dormancy cues than *D. pulex*, which typically inhabits more stable environments. We show that *D. obtusa* invests more heavily in sexual reproduction and resting egg production than *D. pulex* (Fig. 3). Contrary to our hypothesis, both species appear to respond similarly to environmental cues that terminate dormancy (Fig. 5). Together, these findings support the idea that dormancy strategies reflect adaptations to environments that differ in the predictability and degree of environmental change.

The interspecific differences in male and resting egg production we observed are consistent with prior studies examining dormancy across habitats that differ in hydroperiod and environmental predictability (Altermatt & Ebert, 2010; Campillo et al., 2011; Walsh, 2013; Tarazona et al., 2017). Under more variable conditions, selection is expected to favor greater investment in sexual reproduction and resting egg production, even at the cost of reduced short term population growth (Gerber et al., 2018). The increased male and ephippia production that we observed in *D. obtusa* aligns with this expectation, and follows similar patterns reported in other studies of *Daphnia* inhabiting temporary environments (Altermatt & Ebert, 2010). In contrast, populations from more permanent habitats can often persist through extended periods of clonal reproduction, reducing the need for frequent investment in costly sexual reproduction and dormancy (Cáceres & Tessier, 2004). The reduced ephippia production that we observe in *D. pulex* (Fig. 3C) is consistent with a strategy that prioritizes immediate population growth when environmental conditions are relatively predictable. Ephippial fill rates did not differ between species, indicating that once sexual reproduction begins, both species invest similarly in the production of embryos. These results suggest that species differences are more strongly associated with variation in the initiation of sexual reproduction and ephippia production than with differences in hatching outcomes.

The substantial clonal variation we observed within both species also highlights how genetic variation affects dormancy strategies. The intraspecific variation we observe reinforces the idea that dormancy strategies are not fixed within species, and can vary within habitats types (Larsson, 1991; Tessier & Cáceres, 2004). Within a given habitat type, environmental conditions may vary at different spatial and temporal scales, favoring the maintenance of multiple strategies within species (Barnard-Kubow et al., 2022; Porter et al., 2023; Tessier & Cáceres, 2004). Genetic variation in male production and ephippia production suggests that dormancy related traits can evolve in response to local conditions, providing populations with the capacity to respond to changing environments (Barnard-Kubow et al., 2022).

Although early emergence from dormancy was uncommon, it was observed in both species, with no significant differences between species (Fig. 4). As a result, early emergence from dormancy is likely not a dominant strategy in either system, at least under the conditions we provided. However, the presence of low, but non-zero, early hatching is consistent with bet-hedging (Porter et al., 2023). A small fraction of offspring may emerge early to exploit unexpected improvements in habitat quality, instead of undergoing dormancy from which they may not ever emerge. The limited early hatching observed here suggests that this strategy plays a minor role in buffering against uncertainty, rather than serving as a primary mode of dormancy termination.

In contrast to early hatching, late hatching following changes in environmental conditions revealed strong and consistent effects of light, desiccation, and temperature. Across experiments, cold exposure strongly increased the likelihood of hatching, especially when accompanied or followed by exposure to light. These results support a two-step model of dormancy termination, in which cold acts as a permissive cue that relaxes dormancy, while light serves as an activating cue that initiates development (Ślusarczyk et al., 2019). Similar distinctions between cues that terminate dormancy and cues that trigger development have been described in *Daphnia* and other species, and may reflect the need for multiple cues to ensure individuals are emerging into a putatively habitable environment (Finch-Savage & Leubner-Metzger, 2006). The complementary roles of temperature and light also align with differences in cue reliability across habitats. Seasonal temperature declines provide a relatively consistent signal of winter onset in both permanent and ephemeral systems, while increasing photoperiod reliably signals the return of favorable conditions in the spring. The similarity in cue responses between species also suggests that, despite the significant differences in dormancy investment, the environmental requirements for dormancy termination are largely conserved.

One unexpected result was the low rate of hatching after experiencing desiccation treatments. We had hypothesized that individuals from more ephemeral habitats would emerge in response to desiccation as a cue. The inhibitory effect on hatching that we observe may be due to differences between laboratory conditions and the natural environment of ephippia. In the field, ephippia are often covered in sediment or leaf litter that may retain moisture and buffer against complete desiccation (personal observation of ponds in late summer/early fall). Our experimental treatments imposed a harsh desiccation treatment, allowing ephippia to completely desiccate. Alternatively, reduced hatching after desiccation may serve as a survival strategy, where desiccation alone does not trigger emergence and embryos instead require additional environmental cues to terminate dormancy (Pancella & Stross, 1963).

Taken together, our findings support our hypothesis that dormancy strategies are shaped by a balance between environmental predictability and uncertainty. *D. obtusa*, associated with more ephemeral habitats, invests more heavily in the production of dormant offspring. In contrast, *D. pulex*, which is associated with more permanent environments, relies less on dormancy to survive. However, both species retain similar requirements to emerge from dormancy, and both harbor genetic variation in their dormancy strategies across their populations. The presence of both interspecific and intraspecific variation indicates that dormancy strategies vary continuously, likely evolving in response to local environmental conditions (Barnard-Kubow et al., 2022; Tessier & Cáceres, 2004; Walsh & Post, 2012).

## Acknowledgements

The authors thank Dörthe Becker for assistance with field sampling and Heidi Seears for sequencing support. We also thank members of the Bergland lab for helpful discussions and feedback throughout the development of this study.

## Data availability

All scripts and data to perform analyses presented here have been deposited here: https://github.com/rjp5nc/Genoseq_2023/tree/main/DormancyAdaptChap2

Scripts and data will be deposited to DataDryad upon publication.

## Funding

R.J.P. was funded by the CGII Graduate Global Research Grant, the Samuel Miller Fund, NSF EXPAND NRT Fellowship (NSF NRT-ROL 2021791), and the Double Hoo Award (with M.E.D). M.E.D was funded by the University of Virginia College Council Semester award (x2). I.F.M was funded by the University of Virginia Harrison award. A.O.B. was supported by the National Institutes of Health (R35 GM119686) and by start-up funds provided by the University of Virginia.

---

**STable 1:**
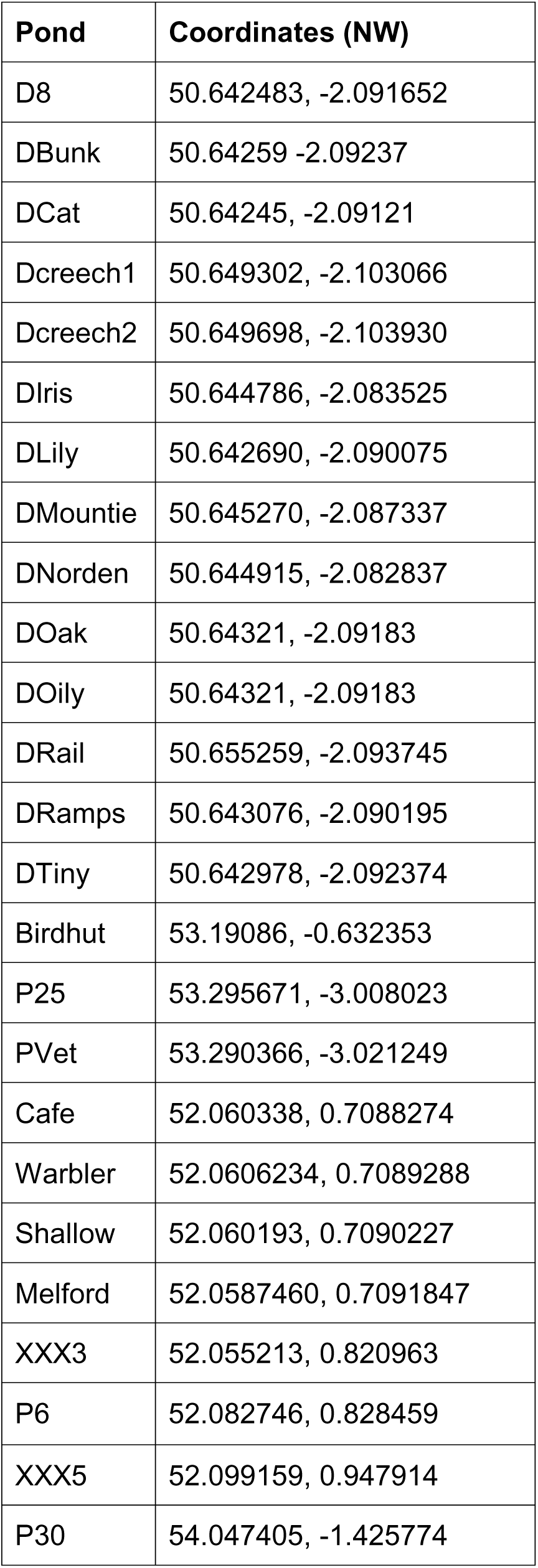
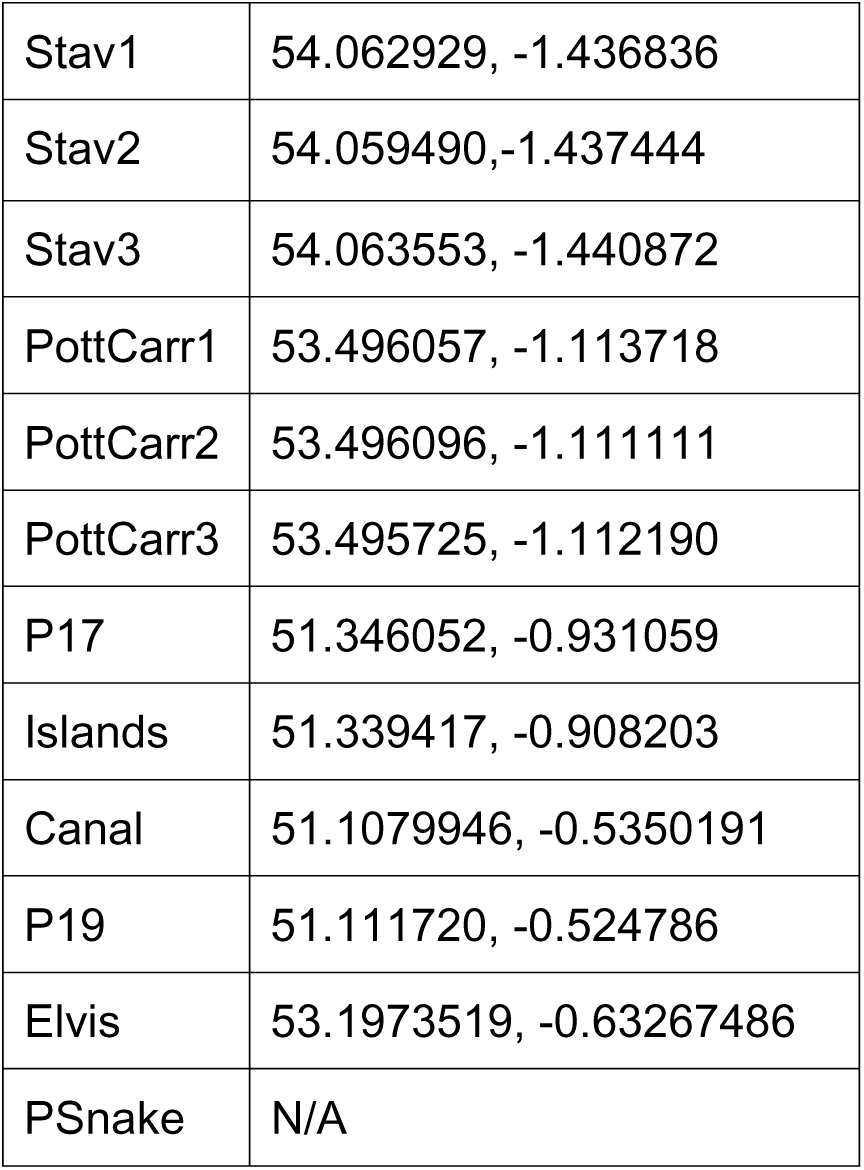
Coordinates of each Pond sampled.

**STable 2:**
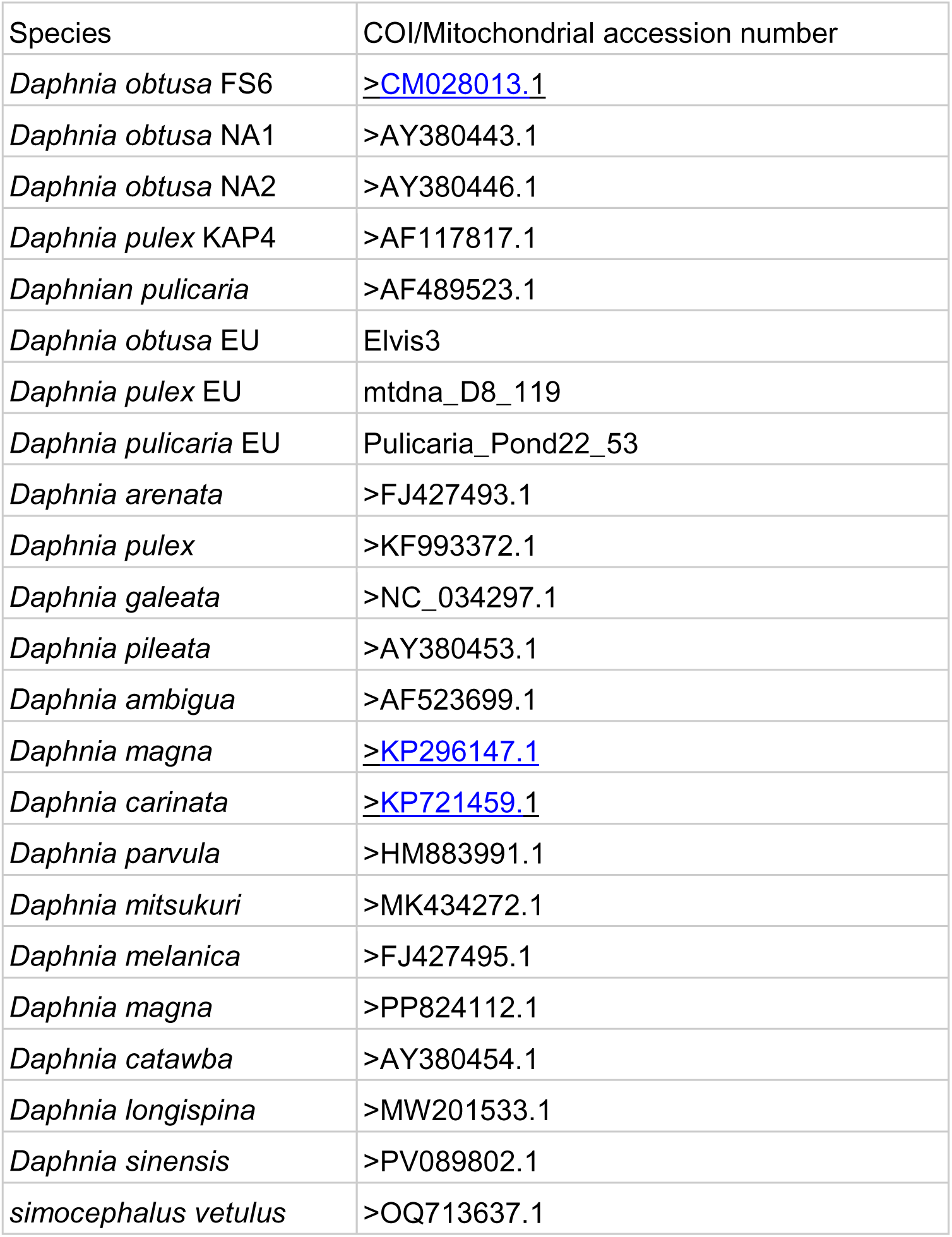
COI reference panel and accession numbers.

| Species | COI/Mitochondrial accession number |
| --- | --- |
| *Daphnia obtusa* FS6 | >CM028013.1 |
| *Daphnia obtusa* NA1 | >AY380443.1 |
| *Daphnia obtusa* NA2 | >AY380446.1 |
| *Daphnia pulex* KAP4 | >AF117817.1 |
| *Daphnian pulicaria* | >AF489523.1 |
| *Daphnia obtusa* EU | Elvis3 |
| *Daphnia pulex* EU | mtdna_D8_119 |
| *Daphnia pulicaria* EU | Pulicaria_Pond22_53 |
| *Daphnia arenata* | >FJ427493.1 |
| *Daphnia pulex* | >KF993372.1 |
| *Daphnia galeata* | >NC_034297.1 |
| *Daphnia pileata* | >AY380453.1 |
| *Daphnia ambigua* | >AF523699.1 |
| *Daphnia magna* | >KP296147.1 |
| *Daphnia carinata* | >KP721459.1 |
| *Daphnia parvula* | >HM883991.1 |
| *Daphnia mitsukuri* | >MK434272.1 |
| *Daphnia melanica* | >FJ427495.1 |
| *Daphnia magna* | >PP824112.1 |
| *Daphnia catawba* | >AY380454.1 |
| *Daphnia longispina* | >MW201533.1 |
| *Daphnia sinensis* | >PV089802.1 |
| *simocephalus vetulus* | >OQ713637.1 |

## Notes

https://github.com/rjp5nc/Genoseq_2023/tree/main/DormancyAdaptChap2

### Competing Interest Statement

The authors have declared no competing interest.

